# Staufen1 localizes to the mitotic spindle and controls the transport of RNA populations to the spindle

**DOI:** 10.1101/2020.03.31.019026

**Authors:** Sami Hassine, Florence Bonnet-Magnaval, Louis-Philip Benoit-Bouvrette, Bellastrid Doran, Mehdi Ghram, Mathieu Bouthillette, Eric Lecuyer, Luc DesGroseillers

## Abstract

Staufen1 (STAU1) is an RNA-binding protein involved in the posttranscriptional regulation of mRNAs. We report that a large fraction of STAU1 localizes to the mitotic spindle in the colorectal cancer HCT116 and in the non-transformed hTERT-RPE1 cells. Spindle-associated STAU1 partly co-localizes with ribosomes and active sites of translation. We mapped the molecular determinant required for STAU1/spindle association within the first 88 N-terminal amino acids, a domain that is not required for RNA binding. Interestingly, transcriptomic analysis of purified mitotic spindles reveals that 1054 mRNAs as well as the precursor ribosomal RNA and lncRNAs and snoRNAs involved in ribonucleoprotein assembly and processing are enriched on spindles compared to cell extracts. STAU1 knockout causes the displacement of the pre-rRNA and of 154 mRNAs coding for proteins involved in actin cytoskeleton organization and cell growth, highlighting a role for STAU1 in mRNA trafficking to spindle. These data demonstrate that STAU1 controls the localization of sub-populations of RNAs during mitosis and suggests a novel role of STAU1 in pre-rRNA maintenance during mitosis, ribogenesis and/or nucleoli reassembly.

**SUMMARY STATEMENT:** Proper localization and functions of macromolecules during cell division are crucial to ensure survival and proliferation of daughter cells.

## INTRODUCTION

The localization of RNA molecules to specific subcellular compartments, a cellular mechanism that is crucial for normal progression of several biological processes, functions to spatio-temporally regulate gene expression (Neriec and Percipalle, 2018; Suter, 2018; Mayya and Duchaine, 2019). Coordination of this post-transcriptional mechanism is controlled by RNA-binding proteins (RBPs) that are thought to bind and regulate overlapping groups of functionally related RNAs (Keene, 2007; Van Nostrand et al., 2020). This mechanism may allow subpopulations of mRNAs to be tagged and functionally grouped into RNA regulons, and ensures that proteins involved in a specific pathway are translated in a highly coordinated fashion.

Staufen1 (STAU1) is a double-stranded RNA binding protein well known for its involvement in the post-transcriptional regulation of gene expression (Wickham et al., 1999; Marion et al., 1999). It is ubiquitously expressed in mammals as alternatively spliced transcripts that generate protein isoforms of 55 kDa (STAU1^55^, STAU1^55i^) and 63 kDa (STAU1^63^) (Wickham et al., 1999; Marion et al., 1999; Duchaine et al., 2000). A large fraction of STAU1-bound mRNAs are associated with translating ribosomes (Ricci et al., 2014; de Lucas et al., 2014; Luo et al., 2002). Genome-wide analyses reveal that STAU1-bound mRNAs code for proteins with heterogeneous functions including transcription, translation, cell growth and regulation of cell cycle (Ricci et al., 2014; de Lucas et al., 2014; Furic et al., 2008; Laver et al., 2013; LeGendre et al., 2013; Sugimoto et al., 2015). Through its binding to specific mRNA populations, STAU1 controls RNA splicing (Ravel-Chapuis et al., 2012), nuclear export (Elbarbary et al., 2013; Ravel-Chapuis et al., 2012), transport and localization (Kiebler et al., 1999; Vessey et al., 2008), translation (Ricci et al., 2014; Dugre-Brisson et al., 2005; de Lucas et al., 2014; Jeong et al., 2019; Sugimoto et al., 2015), and decay (Kim et al., 2005; Kim et al., 2007). STAU1, via the post-transcriptional regulation that it imposes to its bound mRNAs, regulates a wide range of physiologic transcripts and metabolic pathways. STAU1 is crucial for cell differentiation (Kim et al., 2005; Belanger et al., 2003; Gautrey et al., 2008; Yamaguchi et al., 2008; Gong et al., 2009; Kretz, 2013; Cho et al., 2012), dendritic spine morphogenesis (Vessey et al., 2008; Lebeau et al., 2008), long-term synaptic plasticity (Lebeau et al., 2008), a cellular mechanism for long-term memory, response to stress (Thomas et al., 2009), and cell proliferation (Boulay et al., 2014). In addition, misregulation of STAU1-mediated post-transcriptional mechanisms of gene regulation accelerates cancer progression and regulates apoptosis (Xu et al., 2015; Xu et al., 2017; Liu et al., 2017; Damas et al., 2016; Sakurai et al., 2017).

Interestingly, STAU1 expression levels vary during the cell cycle (Boulay et al., 2014). STAU1 levels rapidly decrease as cells transit through mitosis. Its degradation is mediated by the ubiquitin-proteasome system following its association with the E3 ubiquitin ligase anaphase promoting complex/cyclosome (APC/C) via its co-activators CDH1 and CDC20 (Boulay et al., 2014). Therefore, modulation of STAU1 levels by cell cycle effectors may dictate the post-transcriptional expression of its bound transcripts and may contribute to the control of cell proliferation. Accordingly, a moderate overexpression of STAU1 in cancer cells impairs mitosis progression and cell proliferation (Boulay et al., 2014; Wan et al., 2004). Strikingly, STAU1 overexpression has no effect in non-transformed hTERT-RPE1 and IMR90 cells (Boulay et al., 2014) indicating that the types and importance of cellular defects following a modulation of STAU1 levels depends on cellular contexts. Nevertheless, STAU1 is likely to play an important role during mitosis in non-transformed cells as well since its depletion impairs mitosis progression (Ghram et al., submitted).

To understand the role of STAU1 during mitosis, we first documented its subcellular distribution, revealing that an important subpopulation of STAU1 associates with the mitotic spindle. Previous studies have shown that mRNAs can be found on mitotic spindles (Eliscovich et al., 2008; Groisman et al., 2000; Sharp et al., 2011; Blower et al., 2007; Sepulveda et al., 2018; Kingsley et al., 2007; Hussain et al., 2009) but the mechanisms of their transport, localization and post-transcriptional regulation are unclear. We now show that STAU1 is involved in RNA localization on spindle. Using RNA-Seq analysis, we identified RNAs that are enriched on spindles, in particular the 45S pre-rRNA precursor and multiple mRNAs. Interestingly, the pre-rRNA and several mRNAs are delocalized from spindles in HCT116 STAU1-knockout (STAU1-KO) cells compared to wild type (WT) cells. Finally, we show that STAU1 colocalizes with OP-puromycin suggesting that a fraction of STAU1-bound transcripts are locally translated on the mitotic spindles. Altogether, our results suggest that STAU1 regulates the transport/localization of different RNA biotypes and that it may contribute to rRNA maintenance during mitosis and thus to nucleolus reassembly.

## RESULTS

### Localization of STAU1 to the mitotic spindle

To visualize the subcellular localization of STAU1 during mitosis, colorectal cancer HCT116 cells (Fig 1) and non-transformed hTERT-RPE1 cells (Supp Fig S1) were synchronized in late G_2_ by the CDK1 inhibitor RO-3306 and then released from the block with fresh medium. At different time points post-release, cells were solubilized with Triton X100, then fixed and stained with anti-STAU1 and anti-α-tubulin antibodies. DAPI staining was included to visualize DNA. Confocal microscopy analysis of mitotic cells revealed that a significant subpopulation of STAU1 co-localized with α-tubulin on the mitotic spindles (Fig 1 A,B). During all phases of mitosis, STAU1 was observed at the poles of the spindle and also on fibers. During telophase, STAU1 was distributed in the cytoplasm of daughter cells and partly with the remains of polar spindle microtubules. Several controls were included to confirm the specificity of the antibodies (Supp Fig S2).

**Figure 1.**
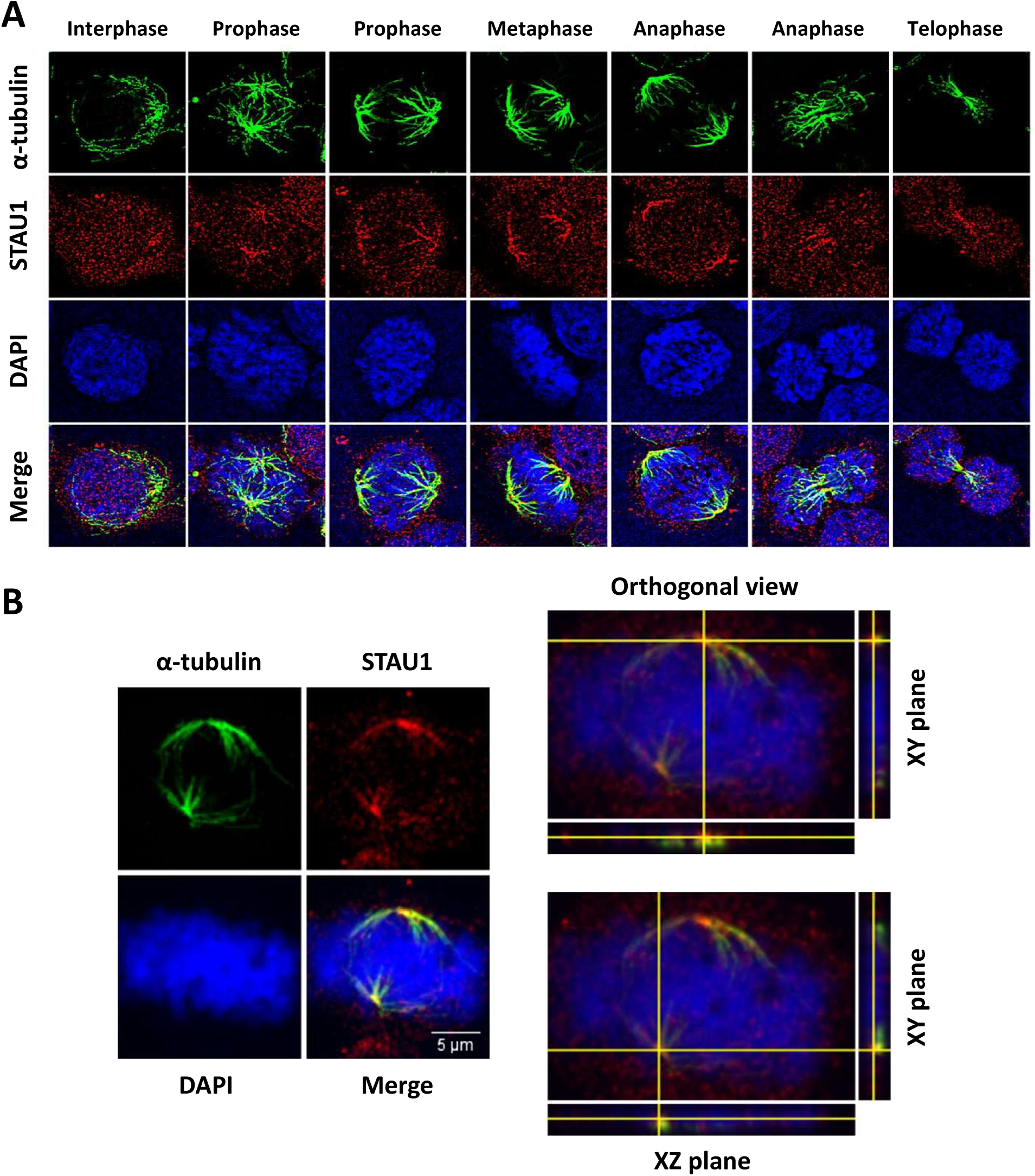
Localization of STAU1 on the mitotic spindle. HCT116 cells were synchronized in late G2 with the CDK1 inhibitor RO-3306 and released from the block to reach mitosis. Mitotic spindle microtubules were stabilized by Taxol. Cells were solubilized with Triton X-100 before fixation to remove soluble material. **A)** Confocal images of cells stained with antibodies against STAU1 and α-tubulin. DNA was stained with DAPI. Cells at different stages of mitosis are shown. This figure is representative of multiple experiments done by three different experimenters. STAU localization on spindle was observed in all mitotic cells. **B)** Left panel: Confocal images of cells stained with antibodies against STAU1 and α-tubulin to visualize STAU1 and mitotic spindles. DNA was stained with DAPI. Right panels: 3D-analysis of the relative position of STAU1 and α-tubulin on microtubules shows co-localization of the proteins in space.

### Biochemical characterization of the mitotic spindle

To confirm a tight association between STAU1 and components of the mitotic spindle, we biochemically purified spindles (Supp Fig S3A) and identified associated proteins by western blotting (Fig 2B; Supp Fig S1B). To prepare spindles, HCT116 and hTERT-RPE1 cells were synchronized in late G_2_ and released. Mitotic cells were incubated with taxol (Fig 2A) to stabilize microtubules and harvested by shake-off. Purified spindles were observed by microscopy to control for the quality of the preparations (Supp Fig S3B). Western blot analysis showed that STAU1^55^ co-purified with tubulin in spindle preparations of both cancer (Fig 2B) and non-transformed (Supp Fig S1B) cells. Interestingly, STAU1^63^ isoform was not detected in the mitotic spindle fraction nor the paralogue protein Staufen2 (STAU2). As expected, aurora A, a known component of the spindle, was found in the spindle fraction while calnexin, β-actin and histone H3 used as negative controls were absent. As a further characterization of the spindle preparations, we showed that the ribosomal proteins S6 (RPS6) and L26 (RPL26) co-purified with spindles, suggesting that both the large and small subunits of the ribosomes, and thus the translation machinery, are present in spindle preparations.

**Figure 2.**
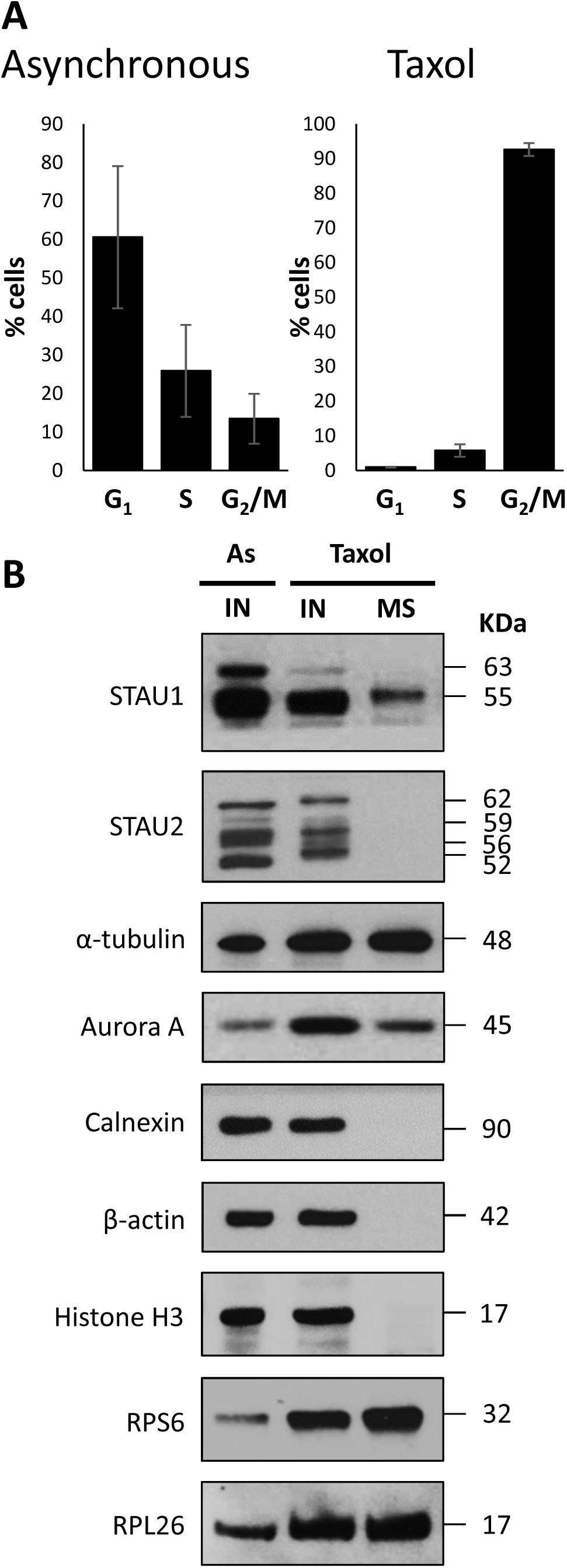
Co-purification of STAU1 with mitotic spindle proteins. **A)** Cell cycle distribution of unsynchronized (Asynchronous) and taxol-synchronized (Taxol) cells was determined by FACS analysis. (*p*-value ≤ 0,05; n = 3). **B)** Western blot analysis of purified mitotic spindles (MS) using specific antibodies. Total input lysates (IN) from asynchronous (As) and taxol-synchronized (Taxol) cells were loaded as controls. This figure is representative of three independently performed experiments.

### dsRBD2 is necessary and sufficient to link STAU1^55^ to spindles

As a first step to define how STAU1^55^ binds to spindle components, we next evaluated the spindle targeting properties of STAU deletion mutants in order to map spindle association determinant. We first generated STAU1-KO HCT116 cell lines (Supp Fig S4A-F) that will prevent putative dimerization between exogenously expressed STAU1^55^ mutants and endogenous STAU1 (Martel et al., 2010). The growth rate of STAU1-KO (clone CR1.3) HCT116 cells was similar to that of WT cells (Supp Fig 4D). We then showed that transiently expressed STAU1^55^-FLAG_3_ co-localized with spindles (Supp Fig S5A) and purified in spindle preparations (Supp Fig S5B) in STAU1-KO cells or in STAU1-control WT cells. Then, STAU1^55^-FLAG_3_ deletion mutants (Fig 3A) were expressed and their presence in the spindle fraction analyzed by western blotting (Fig 3B,C). As shown in figure 3B, mutants that lost RNA-binding activity (3*4*; Δ3) (Luo et al., 2002) were present in the spindle fraction indicating that RNA binding activity is not required for spindle association. These results are consistent with our findings that STAU1^55^ co-purified with spindles even when spindle preparations were treated with RNase prior to Western blotting (Supp Fig S6). Similarly, deletion of the tubulin-binding domain (ΔTBD) had no effect on the interaction with mitotic microtubules. Deletion of RBD4 (Δ4) or RBD5 (Δ5) had no consequence either. In contrast, a deletion that removed the first N-terminal 88 amino acids of STAU1^55^-FLAG_3_, corresponding to RBD2 (Δ2), reduced STAU1^55^ association with mitotic spindles. The reverse experiment in which RBD2-HA_3_ and RBD4-TBD-HA_3_, used as control, were expressed in HCT116 STAU1-KO cells confirmed these results (Fig 4A-C): RBD2-HA_3_ was found in the spindle fraction but not RBD4-TBD-HA_3_. These results indicate that RBD2 is necessary and sufficient for STAU1/spindle association.

**Figure 3.**
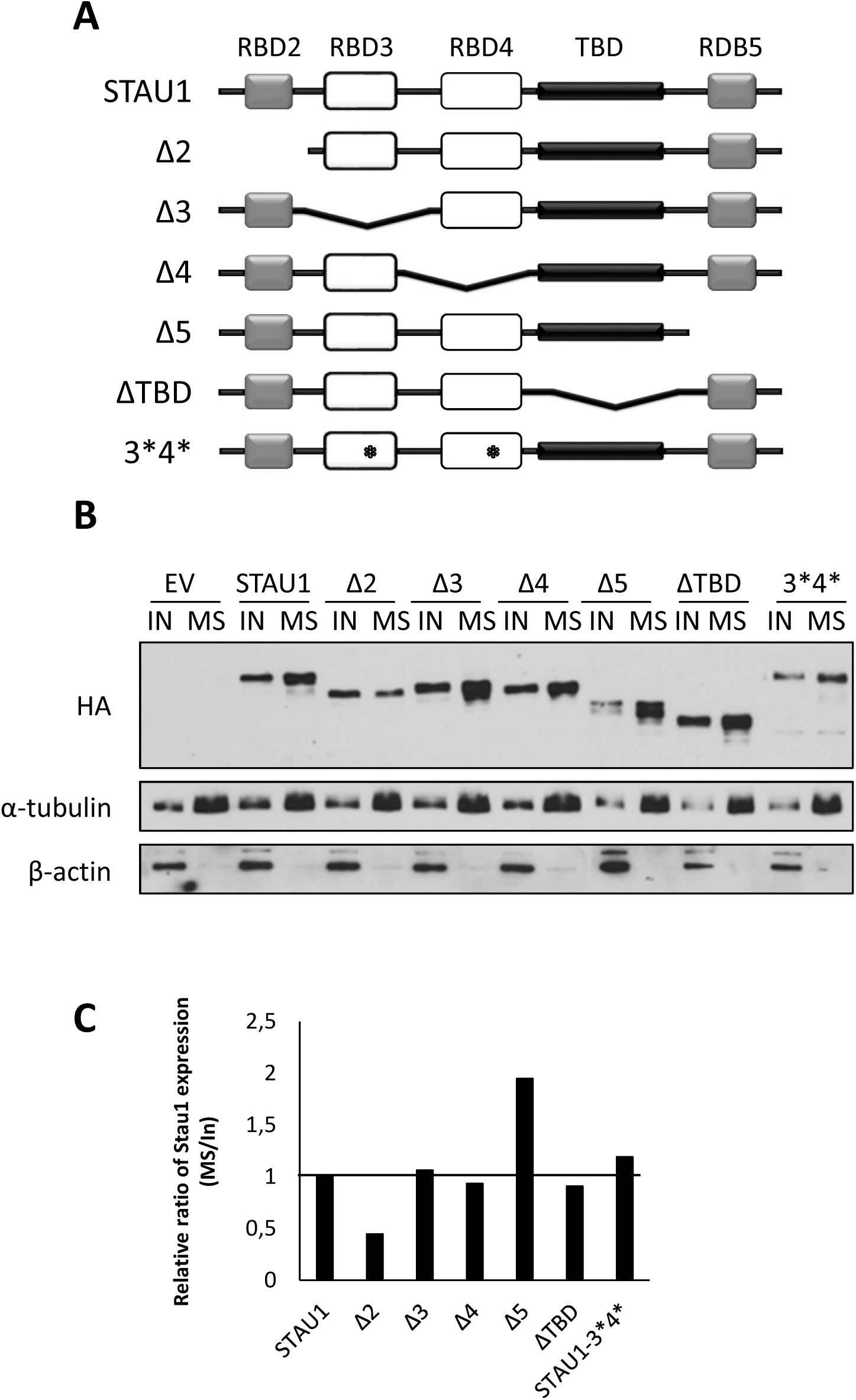
Mapping of the molecular determinant involved in STAU1-spindle association. STAU1-KO HCT116 cells were transfected with plasmids coding for HA_3_-tagged STAU1 WT or deletion mutants (Δ) to map the molecular determinant involved in the binding of STAU1 to the mitotic spindle. **A)** Schematic representation of STAU1 WT and mutants. *, point mutation that abolishes STAU1 RNA-binding activity. RBD: RNA-binding domain. TBD, tubulin-binding domain. White boxes, RNA-binding domains. Grey boxes, domains that do not bind RNA in vitro although they have the consensus sequence for RNA-binding. Black boxes, tubulin-binding domains. **B)** Cells were synchronized in mitosis with taxol. Proteins from the total cell extracts (IN) and from purified spindle preparations (MS) were analyzed by western blotting. Proteins were identified with specific antibodies as indicated. **C)** Quantification of STAU1 proteins on the mitotic spindle. The ratio of STAU1 amounts in the spindle preparations over that in the total mitotic cell extracts was calculated. The ratio obtained for STAU1 WT was arbitrary fixed to 1. These data are representative of two independently performed experiments that gave similar results.

**Figure 4.**
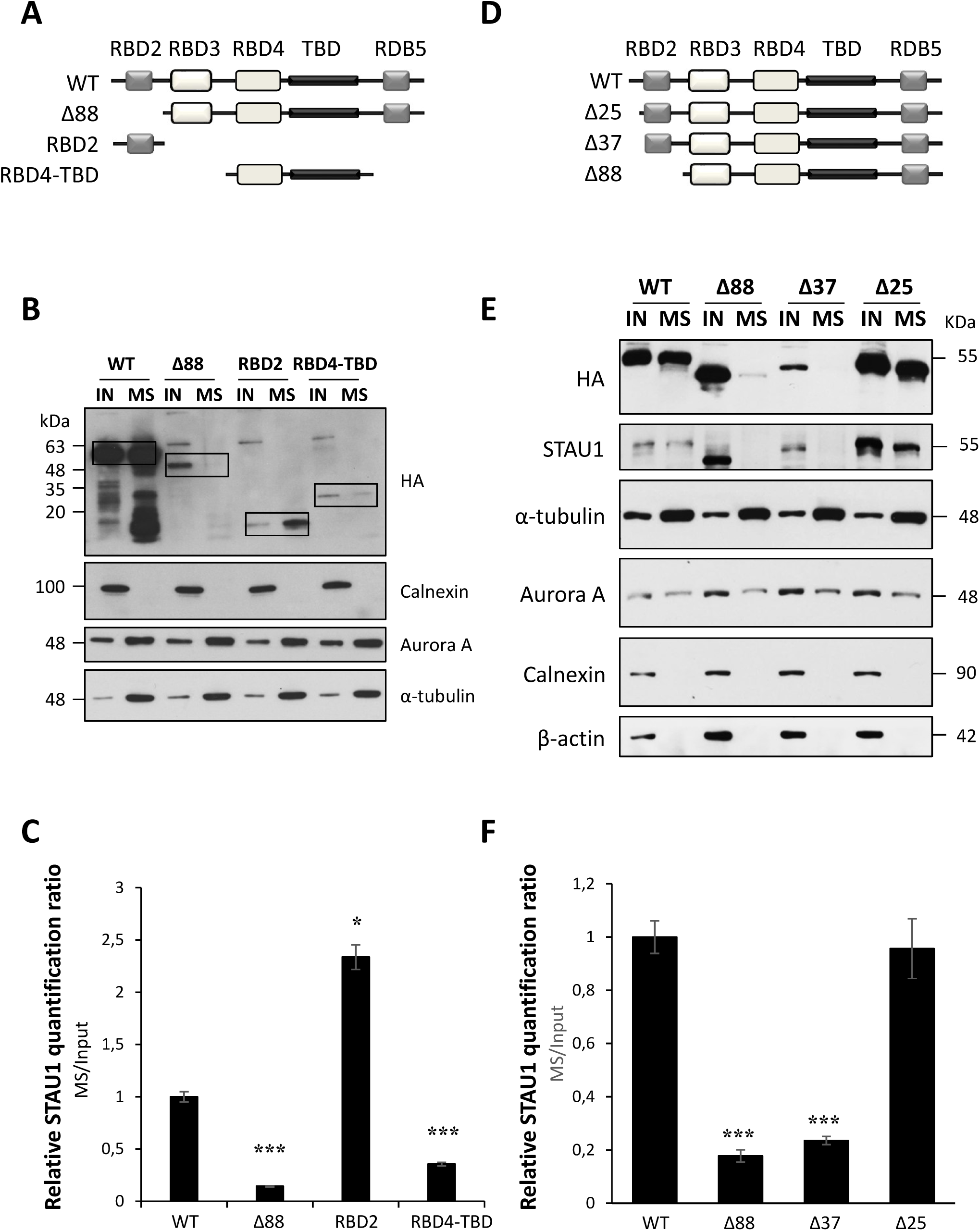
Fine mapping of the N-terminal determinant responsible for STAU1-spindle association. **A, D)** Schematic representations of STAU1 deletion mutants (see legend of figure 3A for box codes). **B, E)** Proteins from total cell extracts (IN) and purified spindle preparations (MS) were analyzed by western blotting with specific antibodies. **C, F)** Quantification of the amounts of STAU1 proteins on the mitotic spindle. The ratio of STAU1 amounts in the spindle preparations over that in the cell extracts was calculated. The graphs represent the means ± s.d. of three independently performed experiments. The ratio obtained for STAU1 WT was arbitrary fixed to 1. *, *p*-value ≤ 0.05; ***, *p*-value ≤ 0.001.

To more finely map the molecular determinant involved in STAU1^55^/spindle association, progressive deletions were made in the N-terminal region and the resulting proteins were tested for their capacity to co-purify with spindles (Fig 4D-F). Western blotting of spindle-associated proteins showed that deletion of the first 25 residues (Δ25) of STAU1^55^ did not prevent STAU1^55^ association with the spindle whereas deletion of the first 37 N-terminal residues (Δ37) abrogated this association. These results indicate that the molecular determinant involved in STAU1^55^/spindle association is located within amino acids 26 and 37.

### Transcriptomes of WT and STAU1-KO cells

Given the well-established and conserved role of STAU1 in the regulation of post-transcriptional gene expression, it is likely that STAU1^55^ is responsible for the transport and/or localization of specific RNAs to the spindle as well as for their post-transcriptional regulation while associated with this structure. Therefore, to highlight a putative role of STAU1^55^ in the transport of RNAs to spindles, we biochemically purified mitotic spindles from parental WT and STAU1-KO cells and identified spindle-associated RNAs by RNA-Seq (Supp Table S1). Total RNAs from WT and STAU1-KO mitotic cells were also sequenced to normalize for putative changes in cell transcriptomes due to STAU1 ablation. RNA Pico chips analysis showed the quality of RNA preparations (Supp Fig S7) and a PCA plot (Khatua et al., 2003) showed that sequencing data are grouped together according to the sources of RNA preparations used (MS or IN) indicating reproducibility of the replicates (Supp Fig 8A). It also indicates that data from whole-cell RNA preparations are different from those from mitotic spindle preparations. Similar conclusions were reached from the calculation of coefficients of correlation between samples (Supp Fig S8B-E).

Comparison of RNA biotypes in total cell extracts (Fig 5A) indicated that the relative expression of RNA types per million reads in the transcriptomes of WT and STAU1-KO cells is similar (Table 1). Almost half of the reads corresponded to protein coding RNAs. Using FPKM of 1 as a threshold value for gene expression, only 108 individual RNAs were found to have altered expression in STAU1-KO cells compared to WT cells (fold change ≥ 2, adjusted p-value ≤ 0.05) (Supp Table S1). 35 protein-coding mRNAs and four lncRNA were upregulated whereas 68 protein-coding mRNAs and one lncRNA were downregulated (Supp Table S2).

**TABLE 1.**
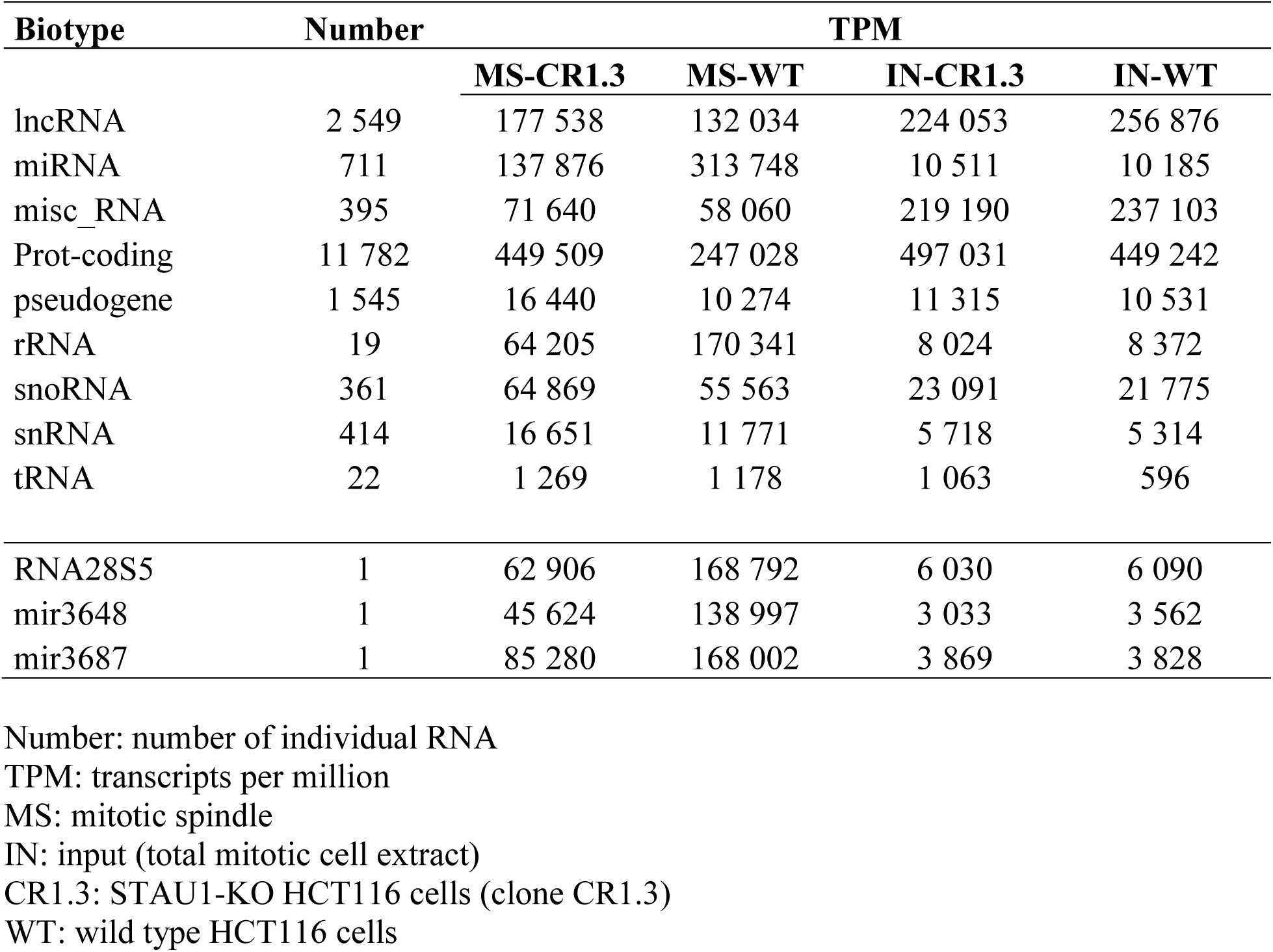
Total number of individual transcripts and of transcripts per million (TPM) across RNA biotypes.

**Figure 5.**
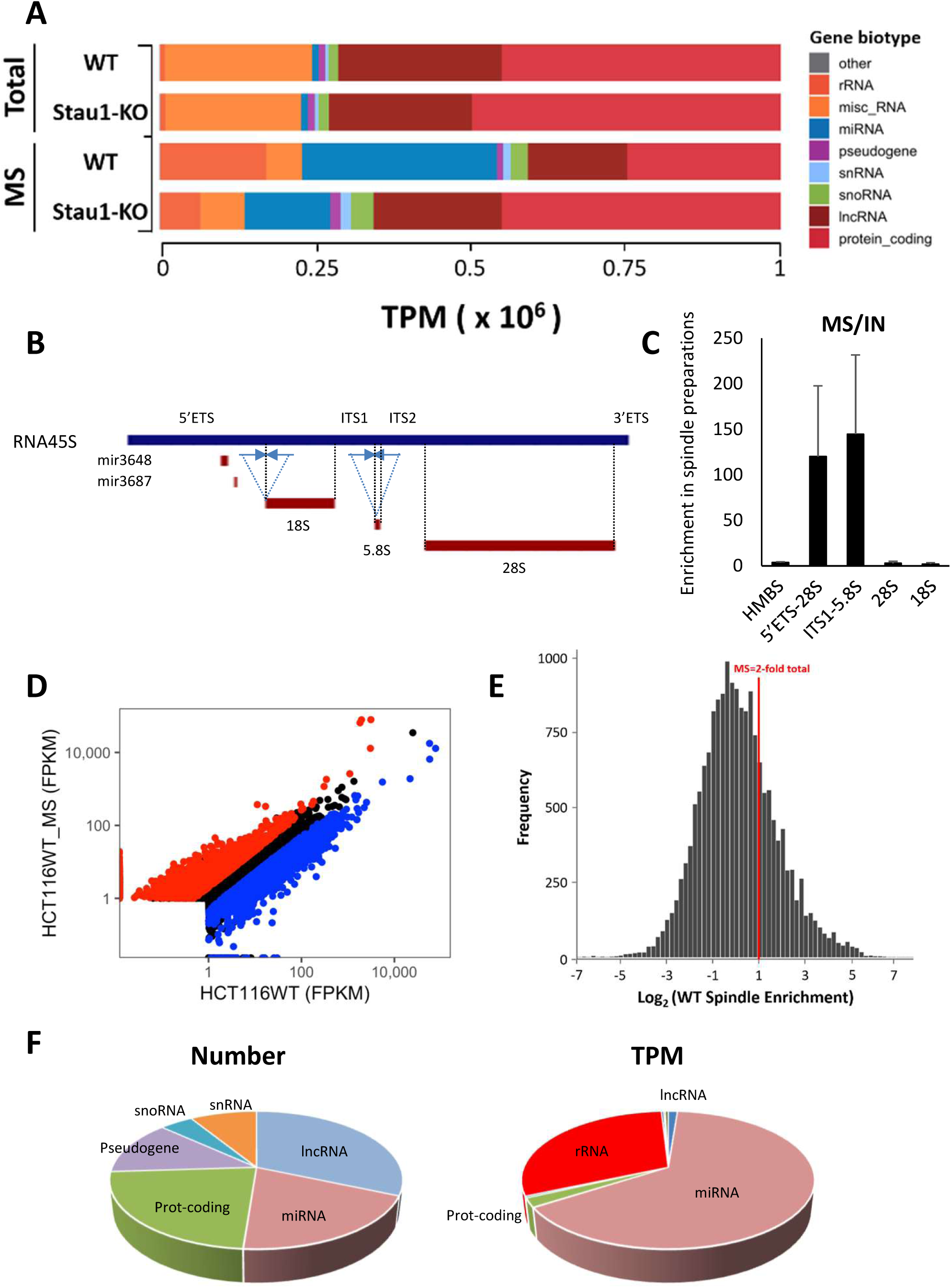
Spindle-enriched transcriptome in HCT116 cells. **A)** Histogram depicting the RNA biotype content, in transcripts per million (TPM), in total mitotic cell extracts (Total) and mitotic spindles (MS) from WT and STAU1-KO HCT116 cells. Biotypes accounting for less than 1% of the overall TPMs were grouped as “other”. **B)** Schematic representation of the RNA28S5 locus showing the 45S pre-rRNA. Arrows indicate the position of the oligonucleotides used for RT-qPCR shown in (C). The position of miRNAs and mature rRNAs is shown. **C)** RT-qPCR amplification of spindle-associated RNAs using oligonucleotides that recognized different parts of the 45S rRNA. The levels of enrichment in spindle preparations (MS) compared to cell extracts (IN) are shown. The graphs represent the mean and standard deviation of three independently performed experiments. HMBS, hydroxymethylbilane synthase; ETS, external transcribed spacer; ITS, internal transcribed spacer. **D)** Scatter plots of the relative distribution of Fragments Per Kilobase of transcript per Million mapped reads (FPKM) in mitotic spindles preparations (MS) vs cell extracts in WT HCT116 cells. Red, RNAs at least 2 fold more abundant (*p*-value ≤ 0.05) in MS than in cell extracts. Blue, RNAs at least 2 fold less abundant (*p*-value ≤ 0.05) in MS than in cell extracts. **E)** Histogram of the frequency of genes in function of their enrichment in MS preparations vs cell extracts in WT cells. **F)** Relative amount of RNAs that are delocalized from the spindle in the absence of STAU1-KO cells. The percentage of the number of different individual RNA in each biotype (left) and the percentage of TPM in each biotype (right) are shown.

### Genome-wide identification of mitotic spindle-enriched RNAs

A different pattern was observed with spindle preparations (Fig 5A. Table 1). The relative expression per million reads of protein coding RNAs was higher in mitotic spindle preparations of STAU1-KO cells compared to WT cells whereas that of miRNAs and rRNAs was lower. Strikingly, the huge decrease of miRNAs and rRNAs is essentially due to three highly abundant RNAs in spindle preparations, RNA28S5, mir3648 and mir3687 (Table 1). Interestingly, the chromosomal location of mir3648 and mir3687 is within the RNA28S5 45S pre-rRNA locus (Fig 5B), suggesting that the two miRNA sequences are present in spindle preparations within the pre-rRNA transcript and not as mature miRNAs. This huge decrease in the number of reads (TPM) of rRNAs and miRNAs in STAU1-KO spindles compared to WT spindles results in a sur-representation of all other RNA biotypes, including protein-coding transcripts, in STAU1-KO spindle (Table 1). To confirm the presence of pre-rRNA in spindle preparations, we RT-qPCR-amplified spindle-associated RNAs with oligonucleotide primers positioned on either side of the 5’ETS/18S and ITS1/5.8S junctions (Fig 5B). Our results indicated that the spacer fragments were linked to ribosomal sequences and therefore that the precursor rRNA is highly enriched in spindle preparations (Fig 5C). Sequences corresponding to mature 18S and 28S rRNAs were also very abundant in the spindle preparations, but they were not enriched compared to input because they are also present as abundant cytoplasmic ribosomes. These results indicate that the precursor 45S rRNA is an important component of the spindle transcriptome.

We then identified other spindle-enriched RNAs in WT cells. We plotted the amount of each RNA (FPKM) in the mitotic spindle preparations in function of their amount in total cell extracts (Fig 5D) and analyzed the frequency of RNAs in function of their enrichment in mitotic spindle preparations compared to cell extracts (Fig 5E). These results identified 1642 RNAs that are enriched at least twice (*p*-value ≤ 0,05) in spindle preparations compared to total cell extracts, including 1054 protein-coding transcripts (Supp Table S3,S4). 28% of these mRNAs are known to bind STAU1 (Furic et al., 2008; Boulay et al., 2014; Ricci et al., 2014; Sugimoto et al., 2015; de Lucas et al., 2014). These mRNAs code for proteins involved in cellular processes such as cell differentiation, GTPase activity, microtubule-based processes, and chromatin organization and modification (Supp Table S5). In addition, the pre-rRNA as well as snoRNAs involved in pre-rRNA processing and MALAT1, a scaffold lncRNA involved in ribonucleoprotein assembly, were highly enriched (Supp Table S3).

### STAU1^55^-mediated localization of RNAs on mitotic spindle

To identify RNAs whose localization to the spindle is dependent on STAU1, we compared the amount of individual RNA in spindle preparations of WT and STAU1-KO cells. We normalized the amount of each RNA in the spindle preparations to that in cell extracts (RNA-spindle/RNA-input) and then compared the ratios in STAU1-KO vs WT cells. We identified 771 individual RNAs, including RNA28S5, miRNAs (including mir3648 and mir3687) and 154 protein-coding mRNAs whose amount in the spindle preparations is at least two-fold lower in STAU1-KO cells than in WT cells (Fig 5F) (Supp Table S6). A different pattern appears with the analysis of TPM: the most important decrease concerns the pre-rRNA and its associated miRNAs while protein-coding transcripts represented only 2% of the total decreased reads (Fig 5F). 29.2% of the protein-coding mRNAs are known to bind STAU1. Among the remaining mRNAs that do not bind STAU1, 60% (60/109) are only marginally expressed (FPKM<2). Bioinformatic analysis (Metascape, A Gene Annotation & Analysis Resources (Tripathi et al., 2015)) indicates that the proteins encoded by these STAU1-bound mRNAs are enriched in GO terms related to regulation of cell shape, actin-cytoskeleton organization, negative regulation of cell growth and differentiation (Supp Table S7).

To confirm the STAU1-mediated differential association of RNAs with spindles of WT and KO cells, selected RNAs were quantified in cell extracts and spindle preparations of WT and STAU1-KO cells by RT-qPCR. Two different STAU1-KO cell lines, generated with different guide RNAs (Supp Fig S4), were used to exclude putative off-target effects. Based on RNA-Seq data, we studied several RNAs whose amounts were decreased in spindle preparations of STAU1-KO cells compared to that of WT cells and are known targets of STAU1 binding. *Hprt* and *rpl22* mRNAs were used as negative controls to normalize data. As expected from RNA-Seq data, the 45S precursor rRNA was delocalized from the spindles of STAU1-KO cells compared to WT cells (Fig 6A) as well as *mex3d, fam101b* and *nat8l* mRNAs (Fig 6B). Used as control, the level of *aspm* RNA was unchanged in STAU1-KO spindle extracts compared to WT extracts.

**Figure 6.**
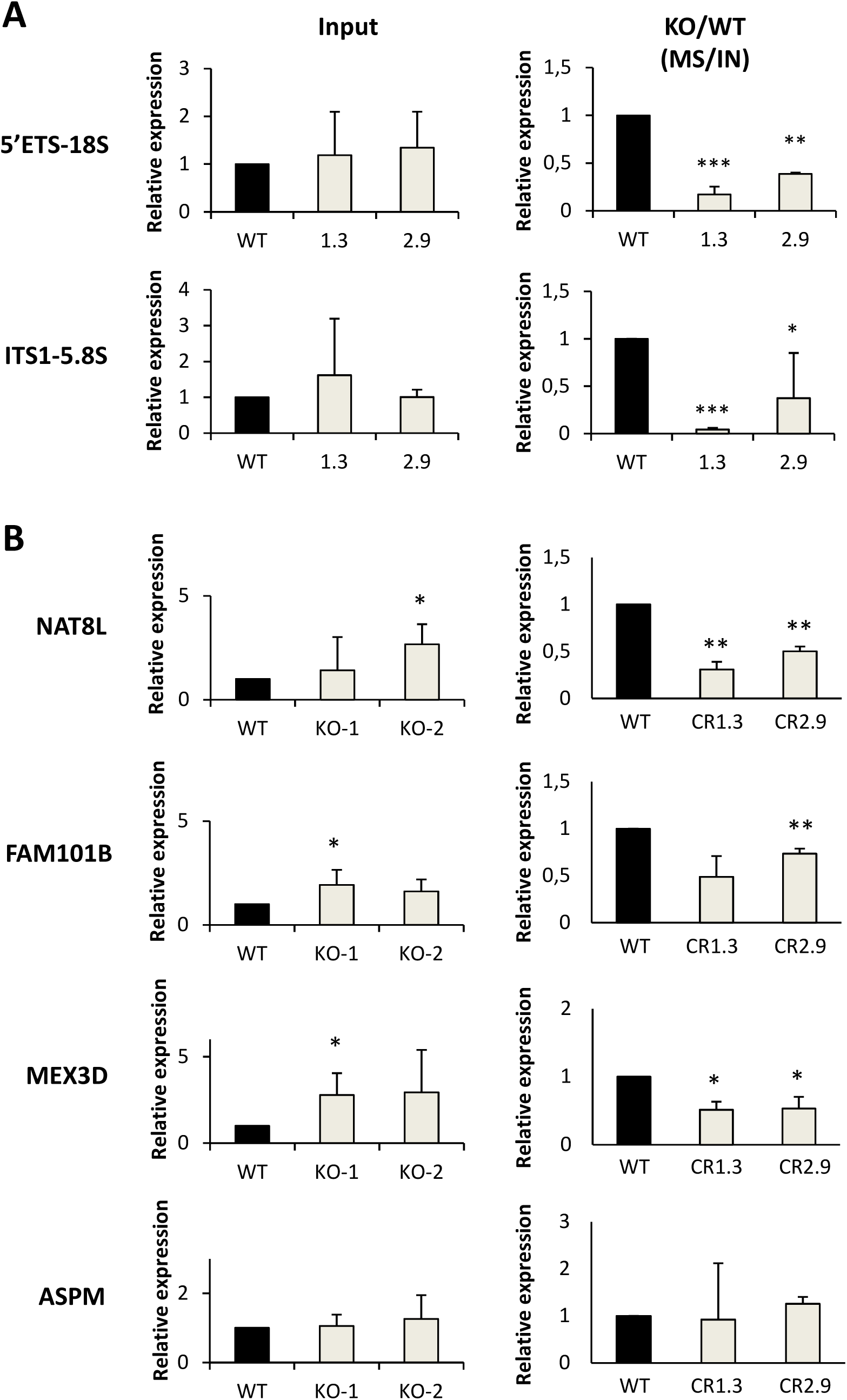
Validation of RNA-Seq data by RT-qPCR. WT and STAU1-KO HCT116 cells were synchronized in mitosis. Two STAU1-CRISPR-KO cell lines (CR1.3 and CR2.9) generated with different RNA guides were used. RNAs in mitotic cell extracts (left) and in purified mitotic spindle preparations (right) were analyzed. RNAs were quantified by RT-qPCR with specific primer pairs. **A)** The contribution of STAU1 in pre-rRNA localization is shown. The graphs represent the mean and standard deviation of three independently performed experiments. *, *p*-value ≤ 0.05; **, *p*-value ≤ 0.01; ***, *p*-value ≤ 0.001. **B)** The amount of mRNAs in cell extracts and in spindle preparations was expressed as the ratio of the amount of a specific gene over that of the negative controls HPRT+RPL22. The ratios obtained with RNAs in the spindle preparations were normalized to the ratio in the cell extracts in both STAU1-KO and WT cells. The ratio in WT cells was arbitrary fixed to 1. Data represents the means and standard deviation of four independently performed experiments. *ASPM*, abnormal spindle microtubule assembly; *MEX3D*, mex-3 RNA-binding family member D; *FAM101B*, refilin B; *NAT8L*, N-acetyltransferase 8 like; *HPRT*, hypoxanthine guanine phosphoribosyl transferase; *RPL22*, ribosomal protein L22.

### STAU1^55^ co-localizes with ribosomes and active sites of translation on the mitotic spindle

The biochemical characterization of spindle-associated proteins indicated that ribosomal proteins co-purified with tubulin and STAU1^55^ in spindle preparations (Fig 2). To study the link between ribosomes, spindles and STAU1^55^, we documented their subcellular localization during mitosis (Fig 7). Using confocal microscopy, we first showed that a significant subpopulation of the ribosomal protein S6 co-localized with tubulin (Fig 7A) and with STAU1 (Fig. 7B) on the mitotic spindle both at the poles and on fibers. Then, we treated cells with O-propargyl-puromycin, a marker of active translation (Chao et al., 2012). The signal was detected by confocal microscopy along with those generated by anti-tubulin and anti-STAU1 antibodies (Fig 7C). Our results indicated that foci of active translation co-localized with subpopulations of both tubulin and STAU1 on the mitotic spindle of individual cells, suggesting that a subpopulation of STAU1-bound mRNAs is locally translated on the spindle while others are not.

**Figure 7.**
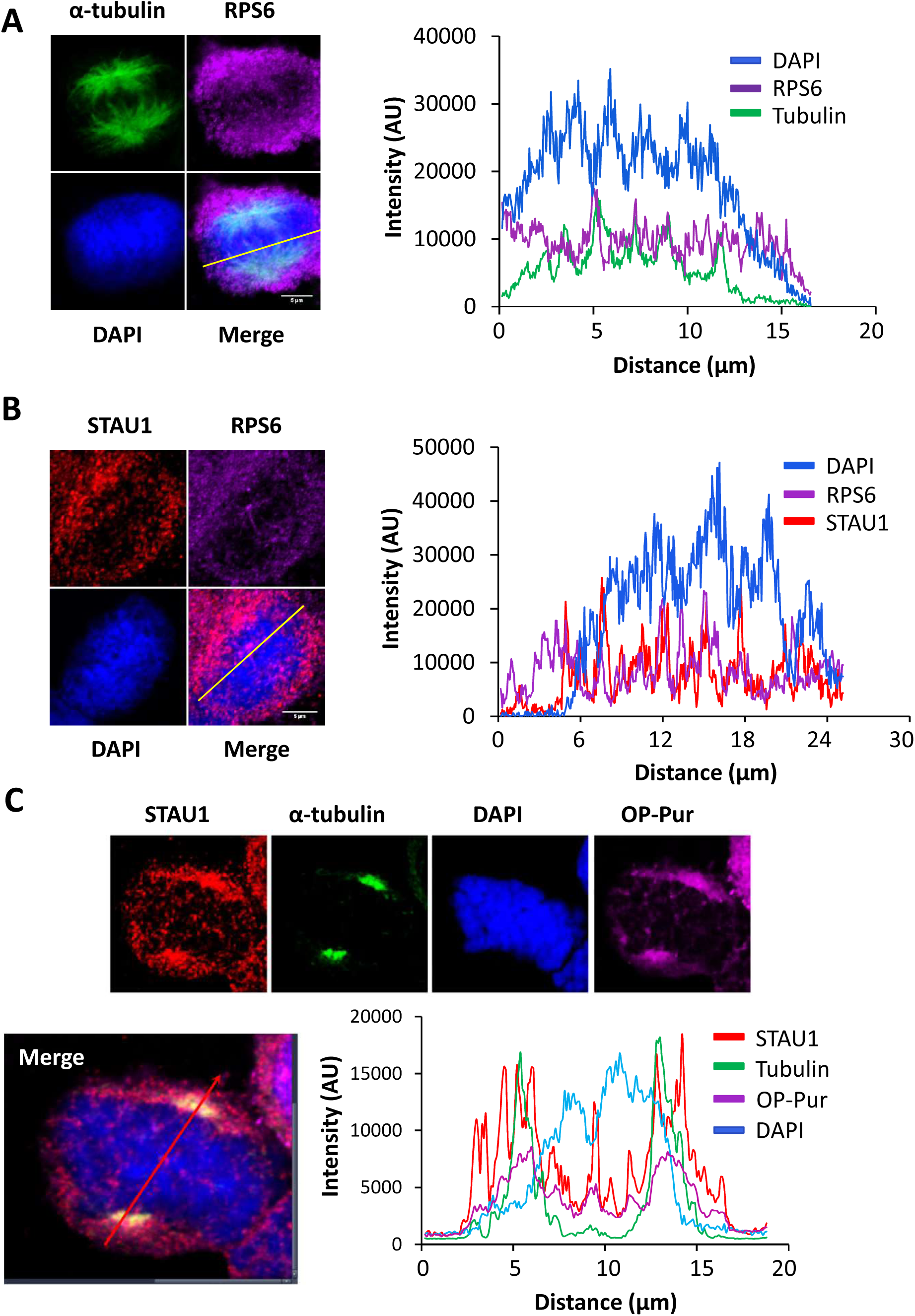
Co-localization of STAU1 and the translation machinery on spindles. HCT116 cells were synchronized in late G_2_ with the Cdk1 inhibitor RO-3306 and released from the block with fresh medium to reach mitosis. Cells were treated with Triton-X 100 to remove soluble materials before fixation. **A)** Cells were stained with anti-S6 (RPS6) and anti-tubulin antibodies to localize ribosomes and the mitotic spindle, respectively. DNA was stained with DAPI. Line scan (right panel) analysis indicates co-localization of RPS6 and tubulin on fiber tracks. The figure is representative of three independently performed experiments giving similar results in every cells. **B)** Cells were stained with anti-S6 (RPS6) and anti-STAU1 antibodies to localize ribosomes and STAU1, respectively. DNA was stained with DAPI. Line scan (right panel) analysis shows overlapping peaks of RPS6 and STAU1 signals. Scale bars = 5 µm. The figure is representative of three independently performed experiments giving similar results in every cells. **C)** Cells were stained with anti-tubulin and anti-STAU1 antibodies as well as with O-propargyl-puromycin (OP-Pur), a marker of active translation. DNA was stained with DAPI. Co-localization was quantified by line scan analysis (bottom). The figure is representative of five independently performed experiments giving similar results in every cells.

## DISCUSSION

In this paper, we show that STAU1^55^ associates with the mitotic spindles in both transformed HCT116 and non-transformed hTERT-RPE1 cells. STAU1^55^ is present in mitotic spindle preparations and co-localizes with tubulin and active sites of translation on the mitotic spindle. This is consistent with previous large scale proteomic studies that identified STAU1 as a spindle component of the human (Rao et al., 2016) and hamster (Bonner et al., 2011) mitotic apparatus. In contrast, STAU1^63^ was not found in spindle preparations. This was unexpected since the sequence of STAU1^55^ is entirely included in that of STAU^63^ (Wickham et al., 1999). It is likely that the additional amino acids at the N-terminus of STAU^55^ to generate STAU1^63^ change the structure of the molecular determinant involved in STAU1^55^ association with the spindle and make it inaccessible for protein interaction. Similarly, the paralog STAU2 is not associated with the spindle, consistent with recent observations that failed to localize STAU2 to mitotic spindle, although it co-localizes with spindle at meiosis I and II (Cao et al., 2016). The human paralogues independently evolved from an ancestor gene to acquire differential biological functions while keeping conserved molecular characteristics. Although the human paralogues are both RNA-binding proteins, they bind mainly different sets of mRNAs (Furic et al., 2008) and are essentially present in distinct ribonucleoprotein complexes (Duchaine et al., 2002). Accordingly, they play different roles in spines morphogenesis (Lebeau et al., 2008; Goetze et al., 2006) and synaptic activity (Lebeau et al., 2008; Lebeau et al., 2011). Interestingly, in drosophila embryo, Staufen protein moves to the pole of the mitotic spindles in close association with the astral microtubules when bicoid 3’UTR mRNA is injected (Ferrandon et al., 1994).

### Molecular determinant involved in STAU1^55^-spindle association

STAU1 is a multi-functional protein with several determinants that control its molecular functions. Notably, BD3 and RBD4 regulate STAU1 RNA-binding activity (Wickham et al., 1999) and RBD4-TBD is involved in ribosome association (Luo et al., 2002). We now show that STAU1 association with spindles requires the N-terminal region that contains RBD2, a domain devoid of RNA-binding activity in vitro (Wickham et al., 1999), although we do not exclude the possibility that RBD2 could bind RNA in vivo as reported for the paralogue protein STAU2 (Heber et al., 2019). This result indicates that STAU1 RNA-binding and ribosome-binding activities are not involved in spindle association. It is interesting to note that deletion of the C-terminal RBD5 facilitates RBD2/spindle association (Fig 3 and 4). This is consistent with previous data showing an interaction between RBD2 and RBD5 (Martel et al., 2010) and indicates that RBD5 may regulate the functions of RBD2. RBD2 was also previously shown to bind CDC20 and CDH1 resulting in STAU1 ubiquitination and degradation during mitosis (Boulay et al., 2014) and to be required for impaired cell proliferation (Boulay et al., 2014). Although mapped in RBD2, the three functions are probably not linked as their molecular determinants do not overlap (unpublished data). In addition, the region of RBD2 involved in STAU1-spindle association (aa 25-37) is also required to increase Pr55^Gag^ multimerization and HIV particle release (Chatel-Chaix et al., 2008) suggesting that HIV Gag may highjack STAU1 function to favor its own replication. Understanding the role of STAU1 on the mitotic spindle may be crucial not only to decipher new pathways leading to cell proliferation but also to discover new steps in RNA virus replication and therefore novel approaches to interfere with them.

The mechanism by which the N-terminal determinant (M^26^RGGAYPPRYFY^37^) allows STAU1^55^ association with spindles is not known. Interestingly, in drosophila, a proline-rich loop in dsRBD2 is required for the microtubule-dependent localization of osk mRNA but not for Staufen association with osk mRNA or for activation of its translation (Micklem et al., 2000). However, mutation of the two prolines (P32A-P33A) did not prevent STAU1 association with spindle (unpublished data), indicating that other amino acids are involved. It will be of interest to test if mutations that prevent tyrosine phosphorylation (Y35 and Y37) or arginine methylation (R27 and R34) (PhosphoSitePlus (Hornbeck et al., 2015)) impair STAU1^55^-spindle association. Alternatively, the N-terminal motif may recruit ubiquitin ligase through two potential ESCRT targeting domains (P^32^PRY and Y^37^PFVPPL) that, in turn, targets the protein to the ESCRT machinery. Interestingly, several ESCRT proteins localize to mitotic microtubules and play important roles throughout mitosis in centrosome localization and duplication, spindle organization and stability, kinetochore attachments, spindle checkpoint, nuclear envelop reassembly and cytokinesis (Dionisio-Vicuna et al., 2018; Morita et al., 2010; Petsalaki et al., 2018; Petsalaki and Zachos, 2018; Vietri et al., 2015).

### Spindle-enriched RNAs

Large scale RNA-Seq experiments identified RNAs that are enriched in spindle preparations compared to total cell extracts, including many protein-coding transcripts (Supp Table S3) (Blower et al., 2007). The fate of these transcripts is not clear. Although it is accepted that essential proteins required for mitosis are synthesized prior to prophase, several studies have shown that mitotic translation (Groisman et al., 2000) and inhibition of cap-dependent translation (Wilker et al., 2007) are important for proper mitotic progression. For example, active translation is needed from late prophase to prometaphase to synthesize proteins that determine the duration of mitosis exit (Cummins et al., 1966). Large scale ribosome profiling experiments confirmed that proteins are synthesized during mitosis (Stumpf et al., 2013; Tanenbaum et al., 2015; Park et al., 2016; Aviner et al., 2013). However, whether translation occurs on the spindle, in the cytoplasm or both is unknown.

The presence of ribosomes and of active sites of translation suggests that translation can occur on spindle, consistent with the presence of several spindle-associated mRNAs in the lists of proteins that are translated during mitosis (Aviner et al., 2013). Mitotic translation indeed contributes to the protein content of the mitotic apparatus (Blower et al., 2007). Consistently, depletion of several of these spindle-enriched mRNAs by RNA interference impairs normal spindle pole organization and γ-tubulin distribution (Sharp et al., 2011), indicating that local translation of these mRNAs on spindle is beneficial for mitosis progression. Interestingly, no correlation was established between the spindle-enrichment of specific mRNAs and their translation on the spindle (Sharp et al., 2011). Therefore, it was proposed that some transcripts are spatially translated on spindles whereas others are translationally-inactive cargos that are later segregated into daughter cells (Blower et al., 2007).

Strikingly, one of the most abundant RNAs on spindle is the RNA28S5 corresponding to the 45S pre-ribosomal RNA. This pre-rRNA is transcribed by RNA polymerase I from multiple 45S rDNA repeat units organized into five clusters on different chromosomes. During interphase, pre-rRNA is found in the nucleolus where it is processed to form mature 18S, 5.8S and 28S rRNAs and assembled into ribosomes (Hernandez-Verdun, 2011). During prophase, the nucleolus is disassembles and rDNA transcription as well as pre-rRNA processing is arrested for the time of mitosis. The 45S pre-ribosomal RNA is maintained during mitosis and was shown to be present in the cytoplasm (Shishova et al., 2011) and at the chromosome periphery together with pre-rRNA processing factors (Sirri et al., 2016; Shishova et al., 2011). During telophase/early G_1_, the nucleolus is reassembled and the mitosis-inherited 45S pre-rRNA is required for regulating the distribution of components to reassembling daughter cell nucleoli (Carron et al., 2012; Dundr et al., 2000). Our study now indicates that the pre-rRNA, as well as numerous snoRNAs involved in rRNA maturation, are associated with the mitotic spindle during mitosis allowing their segregation into the two daughter cells and reassembly of the nucleoli. The presence of the full-length pre-rRNA on spindle contrasts with previous observation in the clam Spisula that only the processed rRNA spacers, but not the full-length precursor, are associated with centrosomes during meiosis (Alliegro and Alliegro, 2013).

### STAU1^55^-dependent localization of mRNAs on the mitotic spindle

Using WT and STAU1-KO cells, we identified RNAs that are delocalized from the spindle when STAU1 is depleted. Since their overall expression in total cell extracts is not changed in STAU1-KO compared to wild-type cells, these results indicate that STAU1^55^ is responsible for the transport/localization of RNA populations to the spindle. Strikingly, the 45S pre-rRNA accounts for most of the reduced reads that are observed in STAU1-depleted cells, revealing a novel role for STAU1^55^ in nucleolus functions and reassembly. STAU1 was previously shown to be associated with ribosomes (Luo et al., 2002) and to enhance translation when bound to the 5’UTR of mRNAs (Dugre-Brisson et al., 2005). In addition, STAU1 was shown to transit through the nucleolus where it is thought to be involved in ribosome and/or ribonucleoprotein biogenesis (Martel et al., 2006). Our results now add an additional putative role of STAU1 in pre-rRNA trafficking during mitosis and nucleolus reassembly in daughter cells.

Trafficking of other RNA populations is also altered in STAU1-KO cells. The number of different protein-coding transcripts represents a relatively small percentage of delocalized RNA molecules (20%) and only 2% of the delocalized reads. We believe that this is an under-representation of the number of delocalized STAU1-bound transcripts due to the incorporation of additional reads required to compensate the huge decrease of rRNA (TPM) in STAU1-KO spindles compared to WT spindles. The fate of these mRNAs on spindle is unknown. It is likely that a subpopulation of mRNAs on spindle is locally transcribed consistent with the colocalization of STAU1^55^ with active sites of translation. Another subpopulation of STAU1-bound mRNAs is likely sequestered to the spindle in a translationally inactive form, and subsequently released and translated during G_1_. Consistently, large scale ribosomal profiling (Stumpf et al., 2013; Tanenbaum et al., 2015; Park et al., 2016) of G_2_, M and G_1_ synchronized cells identified over 300 mRNAs whose translation is up- or down-regulated during mitosis vs G_1_ or G_2_ while the amounts of their corresponding mRNAs remained unchanged. At least 18 of the 154 mRNAs shown to be delocalized from the mitotic spindle in Stau1 KO cells were among those whose translation is regulated. Interestingly, all of them but one show reduced translation during mitosis. Strikingly, 14 of these 18 mRNAs are known targets of STAU1, suggesting that STAU1 may be a critical factor in a mechanism of translation inhibition during mitosis. Finally, we exclude mRNA degradation by the STAU1-mediated decay (SMD) pathway since we previously showed that STAU1-UPF1 association, required for SMD, is inhibited in mitosis (unpublished data).

Thus, STAU1 controls, in different cellular compartments, different sub-populations of pre-rRNAs and mRNAs that likely regulate cell decision during mitosis. Via the post-transcriptional regulation that it imposes to its bound RNAs, STAU1 regulates crucial functions and deregulation of this mechanism may explain the proliferation defects observed in non-transformed cells upon STAU1 depletion (Ghram et al., submitted).

## MATERIAL AND METHODS

### Plasmids and reagents

Plasmids coding for STAU1^Δ2^-HA_3_, STAU1^Δ3^-HA_3_, STAU1^Δ4^-HA_3_, STAU1^Δ5^-HA_3_, STAU1^ΔTBD^-HA_3_, STAU1^3*-4*^-HA_3_, STAU1^Δ25^-HA_3_, STAU1^Δ37^-HA_3_, as well as RBD2-HA_3_ and RBD4-TBD-HA_3_ were previously described (Luo et al., 2002; Martel et al., 2010; Chatel-Chaix et al., 2008; Wickham et al., 1999; Boulay et al., 2014). Monoclonal (1/1000) and rabbit (1/1000) anti-STAU1 antibodies were previously described (Dugre-Brisson et al., 2005; Rao et al., 2019), respectively. Anti-β-actin (A5441, clone AC-74. 1/5000), anti-STAU2 (HPA0191551/500), anti-α-tubulin (T6074, batch number 023M4813. 1/40000), and anti-HA (H6908, batch number 115M4872v. 1/1000) antibodies were purchased from Sigma-Aldrich. Anti-aurora A (30925, batch number 2. 1/2000), anti-α-tubulin (for IF, ab18251, batch number GR201260-1. 1/40000) and anti-histone H3 (ab1791, batch number GR204148-1. 1/3000) antibodies were purchased from Abcam. Anti-calnexin (sc-11397, batch number C1214. 1/1000) antibody was obtained from Santa Cruz. Anti-RPS6 (2212, batch number 4. 1/1000) was purchased from Cell Signaling. Anti-RPL26, (GTX101833. 1/1000) and anti-STAU1 (for IF, GTX106566. 1/200) was obtained from Genetex. Goat polyclonal anti-mouse (p0447, bach number 20051789. 1/3000) and anti-rabbit (p0448, batch number 20017525. 1/5000) antibodies were purchased from Dako.

### Cell culture

The human cell lines hTERT-RPE1 and HCT116 were obtained from ATCC (Manassas, USA). Human colorectal HCT116 cells and STAU1-KO CRISPR-derived clones were cultured in McCoy’s medium with 10% fetal bovine serum, 20 mM glutamine and 1% penicillin–streptomycin (Wisent). The human cell line hTERT-RPE1 was cultured in Dulbecco modified Eagle’s medium (Invitrogen) supplemented with 10% cosmic calf serum (HyClone) or fetal bovine serum (Wisent), 100 µg/ml streptomycin and 100 units/ml penicillin (Wisent). Cells were cultured at 37°C under a 5% CO_2_ atmosphere.

### STAU1-KO HCT116 cell lines

STAU1-KO HCT116 cells were generated by the CRISPR/Cas9 technology (Jinek et al., 2012). Briefly, HCT116 cells were transfected with plasmid coding for GFP, Cas9 and a sgRNA targeting exon 6 of the *STAU1* gene (Horizon Discovery), using Lipofectamine 3000 (Life Technologies/Thermo Fisher). Forty-eight hours post-transfection, GFP positive cells were sorted by FACS and individual cells were grown into 96-well plates until colonies formed. Loss of STAU1 expression was monitored by western blotting using anti-STAU1 antibody. For growth curve assays, cells were harvested every day and the number of cells was counted with an automatic cell counter TC20 (Bio-Rad). The STAU1-KO clone CR1.3 was used in all experiments requiring STAU1 depletion. Clone CXR2.9 was used in the RT-qPCR experiment.

### Cell lysates and immunoblotting

Cells were harvested in phosphate buffered saline (PBS) and lysed in Tris-SDS Buffer (250 mM Tris–HCl pH 7.5, 15 mM EDTA, 0.5% Triton X-100, 5% (w/v) SDS, 100 mM NaCl and 1 mM dithiothreitol) supplemented with a cocktail of protease and phosphatase inhibitors (Sigma-Aldrich) for 10 minutes. Proteins were separated by sodium dodecyl sulfate polyacrylamide gel electrophoresis (SDS-PAGE) and transferred to nitrocellulose membranes (Millipore). Membranes were blocked for 1 hour in PBST (1× PBS, 0.05% Tween 20) containing 5% non-fat dry milk. Primary antibodies were prepared in 1% (w/v) skim milk in PBS-Tween20 (0,2%) and 0,1% (w/v) sodium azide. Membranes were incubated at room temperature with antibodies for one hour or 16 h (anti-STAU2). Secondary antibodies were prepared in 2,5% (w/v) skim milk in PBS-Tween20 (0,2%). Membranes were incubated at room temperature for 1 h with polyclonal goat anti-mouse (Dako) or anti-rabbit (Dako) HRP-conjugated secondary antibodies. Antibody-reactive bands were detected with chemiluminescence substrate ECL kit (GE Healthcare) using ChemiDoc™ MP Systems (Bio-Rad) or X-ray films (Fujifilm).

### OP-puromycin

Active sites of translation were visualized with the Click-iT^®^ Plus OPP Protein Synthesis Assay Kit (Thermofisher) as recommended by the manufacturer. Briefly, cells were incubated with 20 µM Click-iT OPP solution for 30 min, fixed in 3.7% formaldehyde in PBS for 15 min and permeabilized in 0.5% Triton X100 for 15 min. OPP was detected with Click-iT Plus OPP reaction cocktail. Images were acquired with an inverted Axio Observer Z1 confocal spinning disk microscope (ZEISS). Images was processed with the Zen elite blue edition or ImageJ software.

### Immunofluorescence microscopy

Cells were seeded onto poly-L-lysine-treated 20 mm coverslips in a six-well plate at 40% confluence and incubated overnight at 37°C. Cells were permeabilized in 150 mM NaCl, 10 mM Tris (pH 7.7), 0.5% Triton-X-100 (v/v) and 0.1% BSA (w/v) for 5 minutes and then fixed with 4% paraformaldehyde in PBS for 10 minutes at room temperature. Fixed cells were washed thrice in PBS and blocked in PBS containing 0.1% BSA, 0.02% sodium azide and 1% goat serum for one hour at room temperature. Cells were immuno-stained in blocking buffer containing antibodies for 16h at 4°C. Secondary fluorochrome-conjugated antibodies (AlexaFluor 488 goat anti-mouse, AlexaFluor 488 goat anti-rabbit, AlexaFluor 568 goat anti-mouse, or AlexaFluor 568 goat anti-rabbit (Molecular Probes-Invitrogen) were added for 1h at room temperature. Coverslips were washed and mounted on glass slides using ProLongTM Diamond Antifade mountant media (Thermofisher) containing 4’,6-diamidino-2-phenylindole (DAPI). Images were acquired with an inverted Axio Observer Z1 confocal spinning disk microscope (ZEISS). Images processing was performed using Zen elite blue edition or ImageJ software.

### Mitotic spindle preparation

Mitotic spindles were prepared from mitosis synchronized cells essentially as described (Blower et al., 2007). Briefly, mitotic cells were synchronized in late G2 with RO-3306, released and incubated in the presence of taxol (100 μM) for 15 min to stabilize polymerized microtubule. Mitotic cells were collected by shake-off and cell extracts were diluted in lysis buffer (100 mM Pipes, pH 6.8, 1 mM MgSO_4_, and 2 mM EGTA, 4 µg/mL taxol, 2 µg/mL latrunculin B, 0.5% NP-40, 200 µg DNAse 1, 1 U/mL microccocal nuclease, 20 U/mL benzonate, protease and phosphatase inhibitor cocktails) and centrifuged for two min at 700 g. The microtubule pellet was dissolved in hypotonic buffer (1mM PIPES, 5 µg/mL taxol) and centrifuged again at 1 500 g for three minutes to obtain purified mitotic spindle. Microccocal nuclease was omitted when spindle-associated RNAs were purified.

### Genomic DNA sequencing

Genomic DNA was isolated (Bio Basic) and PCR-amplified using the Phusion polymerase (NEB) and specifics primers flanking exon 6 of the *STAU1* gene (sense: 5’ AGCCAAGTTTTTGTCTCAGCC 3’; antisense: 5’ ACAGCTGTCAATGTGCCTTCT 3’). PCR products were cloned into a pBluescript SK (+) vector (Stratagene). Ten clones were randomly chosen and sequenced (Genome Québec).

### RNA extraction and quantitative real-time PCR (qRT-PCR)

Total RNA was extracted using TRIZOL Reagent (Invitrogen) and reverse transcribed into cDNA using the RevertAid™ H Minus Reverse Transcriptase Thermo Scientific™ Kit (Thermo Fisher Scientific). RNA was resuspended in 20 μL water and digested with DNaseI using amplification grade AMPD1 kit (Sigma-Aldrich) prior to reverse transcription. Reverse transcription reactions were done with 1 µg RNA, MuLV RT enzyme and random hexamer (Thermo Fisher Scientific) according to the manufacturer protocol. Resulting cDNAs were qPCR amplified using the Roche LightCycler 480 SYBR Green I Master kit and the LightCycler® 480 instrument (Roche Applied Science). Cycling conditions were set at 95°C for 30 s, 60°C for 30 s and 72°C for 30 s, and 45 cycles. Sense and antisense primers were respectively: ASPM, 5’-GCACCTTTCTGCCATTCTTGAGG-3’ and 5’-TGCTCCACTCTGGGCCATGT-3’; MEX3D, 5’-CAGATGAGCGTGATCGGCA-3’ and 5’-TGTTTGTCTTGGCCCGCAG-3’; FAM101B, 5’-GGCTTTGTCCCCTGTCCTTT-3’ and 5’-GCCTCTCGGAGTCGTACTTG-3’; NAT8L, 5’-CGCTACTACTACAGCCGCAAG-3’ and 5’-CACAATGCCCACCACGTTGC-3; HPRT, 5’-GCTTTCCTTGGTCAGGCAGT-3’ and 5’-CTTCGTGGGGTCCTTTTCACC-3’; RPL22, 5’-TGGTGACCATCGAAAGGAGCAAGA-3’ and 5’-TTGCTGTTAGCAACTACGCGCAAC-3’; ETS-18S, 5’-CGCCGCGCTCTACCTTACC-3’ and 5’-CGAGCGACCAAAGGAACCAT-3’; ITS1-5.8S, 5’-CTCGCCAAATCGACCTCGTA-3’ and 5’-GCAAGTGCGTTCGAAGTGTC-3’.

### RNA-Seq and Differential gene expression analysis

HCT116 were lysed in Trizol reagent (Life technologies) and RNA extracted. TURBO DNA-free™ Kit (Thermo Fisher Scientific) was used to eliminate DNA contamination and RNA was purified with RNeasy mini kit (Qiagen). Ribosomal RNA sequences were removed with the RiboMinus Eukaryote kit for RNA-Seq (ThermoFisher). RNA-Seq libraries were prepared using TruSeq stranded total RNA sample preparation kit (Illumina). Read quality was assessed using FastQC. No trimming was deemed necessary. Read alignment was executed using TopHat on the Human GRCh37 genomes from Ensembl (Trapnell et al., 2012). The GTF annotation file used during the alignment and for counting the number of reads aligned to each feature was also downloaded from Ensembl (release 75). Read count was obtained with featureCounts (Liao et al., 2014). Normalized count values (FPKM) and differential expression was computed with DESeq2 (Anders and Huber, 2010). Gene biotypes and additional information were obtained via the biomaRt R library (Durinck et al., 2009). All correlations and analysis were performed using R. The data discussed in this publication have been deposited in NCBI’s Gene Expression Omnibus (Edgar et al., 2002) and are accessible through GEO Series accession number GSE138441 (https://www.ncbi.nlm.nih.gov/geo/query/acc.cgi?acc=GSE138441).

## ACKNOWLEDGMENTS

We thank Louise Cournoyer in the cell culture facility in the Département de biochimie et médecine moléculaire. We thank the molecular biology and functional genomics core facilities (Odile Neyret, Alexis Blanchet-Cohen) at the Institut de Recherches Cliniques de Montreal (IRCM) and the gene expression analysis core facilities at Genome Quebec Innovation Centre (McGill University) for transcriptomics approaches.

## COMPITING INTEREST

No competing interests declared.

## DATA AVAILABILITY

Supplemental data are available at NAR online.

